# QTrap-Enabled GERD Safety Analysis of Commercial Salsas

**DOI:** 10.64898/2026.07.31.742167

**Authors:** Anastasia Gross, Christopher Singleton, Stefan Gross

## Abstract

Gastroesophageal reflux disease (GERD) is a prevalent chronic disorder where dietary modifications, particularly reducing spicy foods, are a primary management strategy. Salsa, a widely consumed condiment whose spiciness comes from capsaicin, lacks standardized heat labelling, potentially leading to inconsistent capsaicin exposure for consumers. To address this, our study aimed to develop and apply an LC-MS workflow for accurate capsaicin quantification in commercially available salsas. This approach seeks to provide objective “reflux-conscious” spice classification, supporting evidence-based dietary recommendations for individuals with GERD. In the eight commercial brands we examined, we found that products labelled “mild” had significantly lower capsaicin levels as compared to “medium” or “hot”, but that there was an almost 15-fold range of capsaicin within this group. Surprisingly, there was no statistical difference in capsaicin content between those groups labelled “medium” or “hot” facilitating unambiguous assignment to either category, revealing that product labelling alone is insufficient to guide consumers seeking to control capsaicin exposure in their food. The results in this study enable improved brand-specific recommendations for GERD symptom management.

## Introduction

Gastroesophageal reflux disease (GERD) is a common chronic upper gastrointestinal disorder, with meta-analyses and global burden studies estimating a worldwide prevalence of roughly 13–14% and hundreds of millions of affected individuals, making it a major public health issue.[1-3] Pathophysiologically, GERD results from retrograde flow of gastric contents into the esophagus due to a combination of lower esophageal sphincter (LES) dysfunction, transient LES relaxations, hiatal hernia, impaired esophageal clearance, and altered mucosal integrity, often accompanied by peripheral and central sensory hypersensitivity. [4-6] Current international guidelines identify proton pump inhibitors (PPIs) as a primary front-line therapy because of their proven efficacy in healing erosive esophagitis and controlling heartburn.[7-9] Alongside pharmacologic therapy, dietary modification— including reduction of spicy foods—has also been demonstrated in clinical and observational studies to be an effective front-line strategy for reducing GERD symptom burden, with structured dietary interventions consistently improving heartburn frequency and severity. [10-12]

Salsa has become one of the most widely consumed condiments in the United States, with market analyses showing that salsa sales have surpassed those of traditional condiments such as ketchup, making it a major dietary source of spicy foods.[13] The characteristic heat of salsa is largely attributable to capsaicin, the principal pungent capsaicinoid found in chili peppers such as *Capsicum annuum* (e.g., jalapeño, serrano) and *Capsicum chinense* (e.g., habanero), which contribute substantially to the sensory intensity of commercial salsas. [14, 15] Spiciness is commonly expressed in Scoville Heat Units (SHU), a scale originally based on the Scoville organoleptic test, in which trained tasters evaluated serial dilutions of pepper extracts; although widely used, this method is inherently subjective due to individual variability in sensory thresholds. [14] Modern analytical approaches quantify capsaicinoids using high-performance liquid chromatography (HPLC), but the Scoville scale itself remains rooted in sensory perception and lacks cross-product standardization.[15, 16] Importantly, the U.S. Food and Drug Administration (FDA) does not regulate or define “mild,” “medium,” or “hot” heat levels for salsa or other spicy foods; FDA regulations instead focus classify salsa as an acidified food on pH requirements for acidified foods, not spiciness.[17] Consequently, although manufacturers routinely label salsa with heat descriptors, these labels are not based on any universal or federally standardized metric.

Recent advances in analytical chemistry have enabled highly sensitive quantification of dietary irritants such as capsaicin and caffeine using liquid chromatography-mass spectrometry coupled approaches. [18-21] Quantitative LC-electrospray ionization mass-spectroscopy methods have demonstrated detection limits in the picogram-to nanogram range with excellent linearity and recovery characteristics for capsaicin and related compounds. [22] The analytical specificity of LC-MS therefore offers a promising objective assessment of capsaicin burden in commercial food products.

Because dietary management remains a cornerstone of symptom control in patients with GERD [11, 23], quantitative characterisation of irritant exposure may provide clinically meaningful dietary guidance. Capsaicin activates transient receptor potential vanilloid (TRPV1) receptors located throughout the gastrointestinal tract, producing sensations of burning and irritation that may worsen reflux-associated discomfort in sensitive individuals.[24-26] Although spicy foods are commonly discouraged in GERD dietary recommendations, little standardization exists regarding the actual capsaicinoid content of prepared commercial foods. Consequently, consumers relying on qualitative descriptors such as “hot” or “medium” may unknowingly ingest capsaicin concentrations substantially different from expected levels. We therefore hypothesized that commercially available salsas could exhibit marked variability in capsaicin concentration independent of product labelling, and that LC-MS-based quantification could serve as the basis for a more objective “reflux-conscious” spice classification system. Such an approach could bridge analytical food chemistry and preventative dietary medicine by translating chemical measurements into practical consumer guidance.

The objective of the present study was to develop and apply an LC-MS workflow for quantifying capsaicin concentrations in a representative panel of commercially available salsa products. We further sought to compare experimentally measured capsaicin abundance with manufacturer-designated heat classifications to evaluate the consistency of consumer-facing labelling practices. Given the demonstrated sensitivity and robustness of LC-MS methods for capsaicinoid analysis, this platform provides a suitable analytical basis for standardized assessment of pungency exposure in complex food products. By establishing quantitative capsaicin profiles across commonly consumed salsas, this work aims to support the development of evidence-based dietary recommendations for individuals susceptible to reflux symptoms and to illustrate the broader utility of analytical chemistry in preventive nutrition and food safety applications

## Methods

### Sample Preparation

Commercial salsas as described were obtained from supermarkets in the Boston, Massachusetts area. To ensure homogeneity, bottles were shaken at 2500 RPM in a Benchmark Scientific BV1010-TST for 10 minutes before transferring15 g into a 50 ml Falcon tube. To each tube, 1 g of glass beads (Sigma, G1277) was added and the samples were homogenized for 30 minutes at room temperature at 2500 RPM, then centrifuged for clarification at 5000g for 30 minutes. Deproteination was accomplished by withdrawing 500 ul and mixing with an equal volume of ice-cold methanol containing 100 ng/ml buspirone (Sigma, B7148) internal standard followed by centrifugation at 20,000g for 30 minutes at 4°C. Pure capsaicin standard (Sigma, M2028) was prepared as a 1 mg/ml stock solution in 100% methanol. All salsa samples were prepared and analysed in triplicate.

### Mass spectrometry

Analysis was performed on a Sciex5500 QTRAP operating in positive MRM mode. The LC inlet was a Waters Acquity system, consisting of a Binary Solvent Manager pump, Sample Manager autosampler, and Column Manager. Mobile phase A was 10mM ammonium formate with 0.1% formic acid in water, and mobile phase B was acetonitrile with 0.1% formic acid. The gradient program was 95:5 mobile phase A:mobile phase B from 0 to 0.6 minutes, with a ramp to 95% B by 1.65 minutes, a hold until 2.1 minutes, and a return to initial conditions of 95:5 A:B at 2.2 minutes. These initial conditions were held until 3 minutes, at which point the system prepared for the subsequent injection. The main MRM transition for the quantitation of capsaicin was 306.2->137.0, with confirmatory qualitative transitions of 306.2->122.0 and 306.2->107.0. The collision energy (CE) for the transitions was 30, 60 and 40, for the fragments 137.0, 122.0 and 107.0, respectively. The internal standard buspirone was monitored with an MRM transition of 386.0->222.0, using a collision energy of 30. For all ions monitored, the common parameters are as follows: Curtain gas 35, Ionspray voltage 5500, Temperature 550, Gas 1 60, Gas 2 60, CAD gas high, Entrance potential 10, and collision exit potential 15.

Chromatography was performed in low pH with acetonitrile as the mobile phase modifier on a Acquity CORTECS C18 1.7uM, 2.1x50mm column at a flow rate of 0.650 ml/min with the following parameters: Solvent A: 10mM Ammonium formate + 0.1% formic acid; Solvent B: Acetonitrile + 0.1% formic acid; Sample Injection Volume: 15uL Column temperature 50°C. The capsaicin standard curve was prepared from the initial stock solution, ranging from 2000 ng/ml to 1 ng/ml and showed linearity of 0.9 over the entire range.

## Results and Discussion

Through the use of an extremely sensitive mass spectrometer, this study enables a simple dilute-and-shoot method for LC-MS sample preparation that significantly speeds analysis compared to other methods which require extensive sample handling like solid-phase or liquid-liquid extraction. [27-30] This allowed for the processing in parallel of a large number of samples and replicates, facilitating a broad sampling of the labelled spice levels and brands available to the average consumer. We present the direct measurement of capsaicin content for eight different brands across the three generally accepted descriptions of spice levels: “mild”, “medium”, and “hot”. In addition, the Taco Bell brand also sells a product with an even higher level of spice, “diablo”, which were included in our analysis.

The capsaicin content of the eight brands we measured is presented in Fig 1A. The average and standard deviation measured capsaicin content for “mild” products was 5.8 ± 5.1 µg/g of salsa. For “medium” products, the measured content was an average of 64.8 ± 47.6; and for “hot” products, 114.4 ± 87.8. The precision of the assay is quite high, with the relative standard deviation for the measurement of capsaicin for an individual sample being uniformly less than 10%.

**Fig 1.**
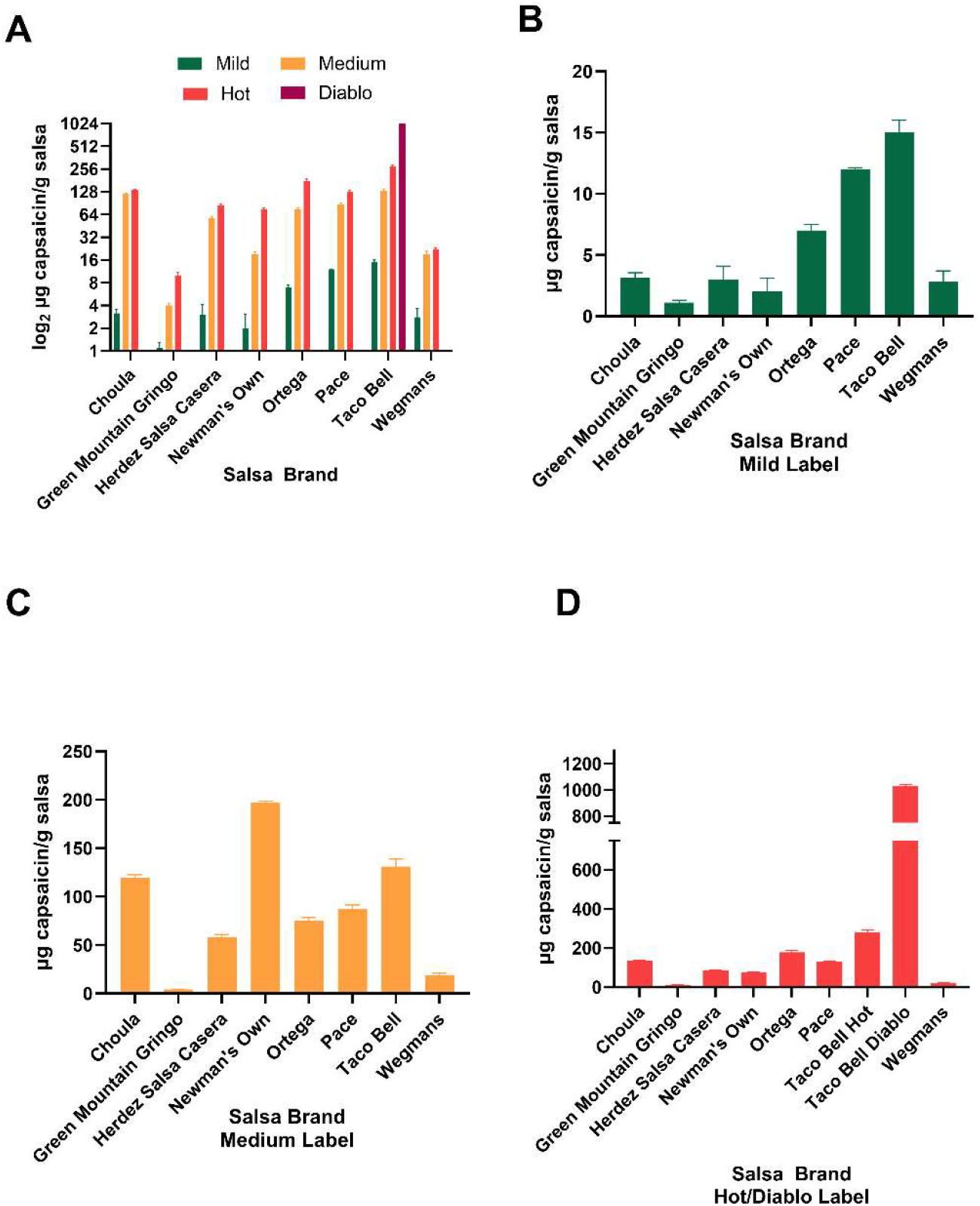
Capsasin content of commercial salsa broken down by brand and label claim of spice levels. (A) A general overview of the measured capsaicin content of the materials in this study. Bar height represents the average capsaicin content averaged over three independent sample preparations, with the error bars designating the standard deviation of the mean. Y-axis has been log_2_-scaled to facilitate accommodation of all the measurements on a single graph. (B-D) Capsaicin content of salsas broken down by label claim; “mild”, “medium”, “hot”; the Taco Bell-claimed extra spicy “diablo” is included in the final figure.

For GERD-suffering consumers seeking to minimize their capsaicin consumption our study would suggest that the Green Mountain Gringo “mild”, with a capsaicin content of 1.1 ± 0.2 ug/g salsa would be the best choice. Should that be unavailable, the Newman’s Own “mild” with a capsaicin content of 2.0 ± 1.1 ug/g salsa would be a possible second choice. The Taco Bell “mild” brand, with a measured content of 15.0 ± 1.0 ug/g capsaicin should be avoided, as it has a capsaicin content almost 15 times as high as Green Mountain Gringo “mild” despite bearing the same “mild” label description for capsaicin content. Overall, the Green Mountain Salsa brand had the lowest content of capsaicin in every label category, while Taco Bell had the highest. Capsaicin content by brand and label is broken out in Fig 1B through 1D).

For consumers willing or able to consume higher levels of capsaicin in the interest of flavor enhancement, the picture is considerably more cloudy when seeking to select a brand and spice-level label claim with specific intention. In Fig 2, we present an analysis of our data aggregated by label claim. We used Student’s T-test[31] to compare the overall populations between the three label claims. Despite the individual differences described above, overall the salsas in the “mild” category as whole had a statistically significant difference in capsaicin content as compared to the “medium”, p = 0.010, and the “hot”, p = 0.010. However, the difference in the means of measured capsaicin content between the “medium” and the “hot” categories, given the large degree of variation in capsaicin content between individual brands was not significant, with a p = 0.182. This suggests that in a random sampling of brands, at the higher two levels of label claim of spicyness, the actual label claims would have a poor chance of matching the actual capsaicin content of the product. For example, the Taco Bell “medium” with a capsaicin content of 131.2 ± 7.6 actually exceeds the Green Mountain Gringo “hot” which has a capsaicin content of 10.2 ± 1.1 ug/g – a difference of more than 12-fold. We did find, however, that within individual brands the overall trend of label claim matching capsaicin content was uniformly preserved.

**Fig 2.**
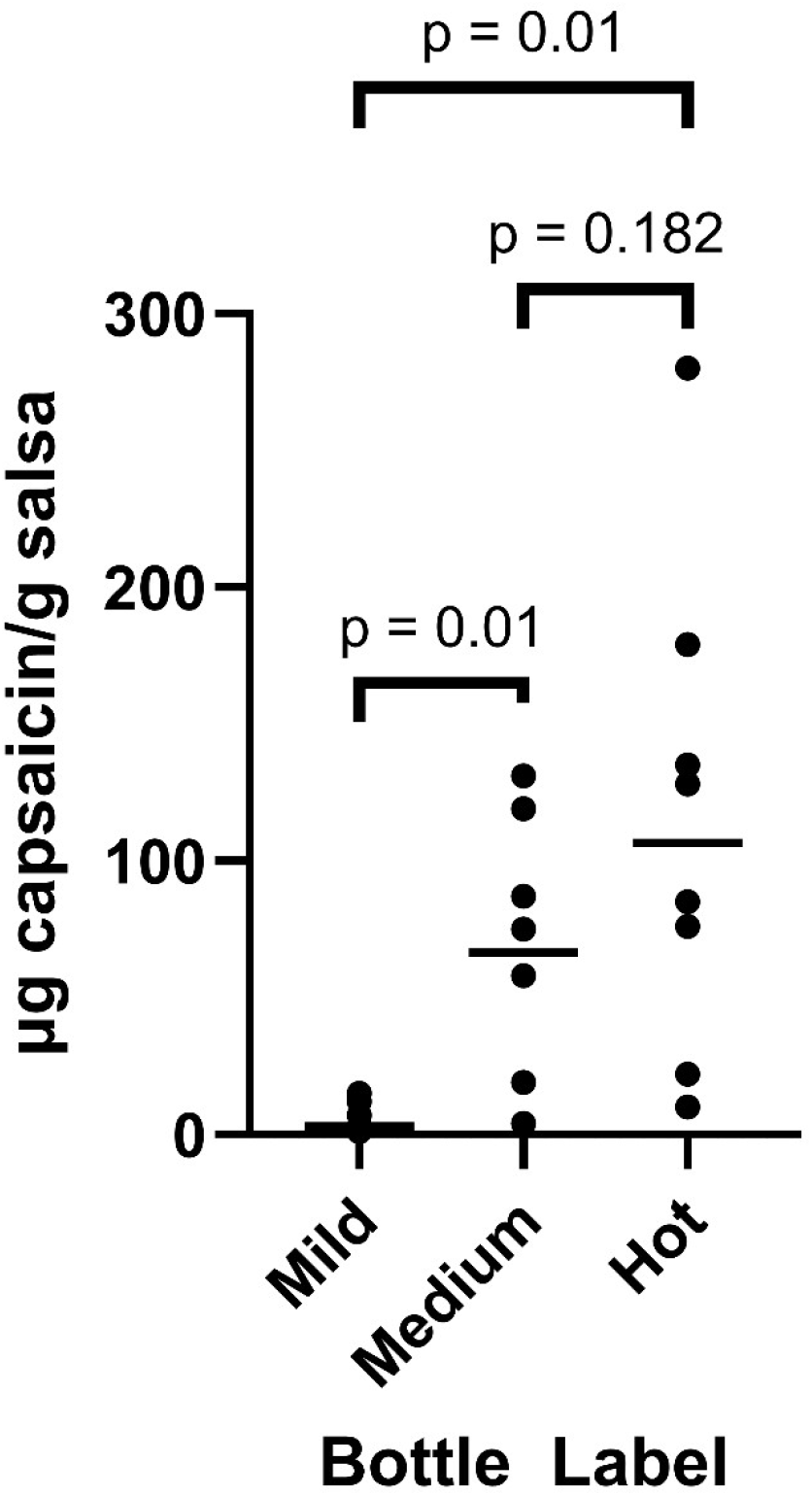
Capsaicin content of salsas by label claim. Each dot represents the average of the measured capsaicin content of three replicate salsa preparations. Short horizontal lines indicate the mean content per group. Student’s t-test p values for comparison between the indicated groups are indicated by brackets. The Taco Bell “diablo” as a unique label specific to the brand was not included in the analysis.

Finally, for consumers seeking a high level of capsaicin content in their food, the three highest capsaicin-containing salsas were in Ortega’s 179.2 ± 9.8 , and Taco Bell’s “hot” and “diablo” at 280.1 ± 11.3 and 1029 ± 14.6 ug/g respectively. The Taco Bell “diablo” implies an extreme level of capsaicin content with the marketing implication that Taco Bell’s “diablo” label contained a product level more spicy than competitors’ “hot”, and thus differentiated and more attractive to consumers seeking very spicy food. We found this implication to be valid, although as the “diablo” label was unique in our sample set, there was no possible comparison to other brands’ label claims.

Our study revealed a wide range of capsaicin content between brands, even when all brands adhere to a common three-level “mild”. “medium”, and “hot” labelling scheme. We have identified two brands of salsa most suitable for GERD-suffering patients seeking improve their symptoms by pursuing a low-capsaicin diet while preserving flavor, variety, or ethnic food identity. While the tripartite labelling scheme was a suitable guidance framework within the same brand, our data suggests that utilizing the same strategy would not be effective when comparing between brands. Further extensions of this study could involve sampling a greater number of salsa brands as well as examining batch-to-batch variations between identically labelled products. This would be particularly interesting because a number of environmental and manufacturing factors, such as metabolic adaptations to growing conditions, location of the capsaicin fruit on the plant, and post-harvesting handling can result in changes in capsaicin levels in the final product. [32,33] We also believe that our study reveals shortcomings in the regulatory framework around salsa production and labelling and suggests consumers, especially GERD patients, would be better served with a regulatory policy extension to 21 CFR 101.4 requiring the disclosure of capsaicin content in foods.

## Acknowledgments

The authors wish to recognize Mr. Jason McLean, Head of Science, at the British International School of Boston for scientific pedagogy; and Mr. William Warren, Senior Scientst, Bioanalytical Mass Specroscopy, of Servier Pharmaceuticals for helpful technical discussions.

